# A novel bacteriocin produced by *Bifidobacterium longum* subsp. *infantis* has dual antimicrobial and immunomodulatory activity

**DOI:** 10.1101/2022.01.27.477972

**Authors:** Sree Gowrinadh Javvadi, Magdalena Kujawska, Diana Papp, Aleksander M Gontarczyk, Anne Jordan, Melissa A.E. Lawson, Ian J. O’Neill, Cristina Alcon-Giner, Raymond Kiu, Paul Clarke, Naiara Beraza, Lindsay J Hall

**Affiliations:** Gut Microbes & Health, Quadram Institute Bioscience, Norwich Research Park, Norwich, NR4 7UQ, UK; Intestinal Microbiome, School of Life Sciences, ZIEL – Institute for Food & Health, Technical University of Munich, Freising, 80333, Germany; Norfolk and Norwich University Hospitals NHS Foundation Trust, Norwich, NR4 7UY, United Kingdom; Norwich Medical School, University of East Anglia, Norwich Research Park, Norwich NR4 7TJ, United Kingdom

## Abstract

Bacteriocins are ribosomally-synthesized antimicrobial peptides produced by bacteria with either narrow or broad spectrum activity. Many genome mining studies have indicated that bacteriocin gene clusters are widespread within certain gut microbiota members. In early life, *Bifidobacterium* comprise the dominant microbiota genus in vaginally delivered and breast-fed infants, with high levels associated with improved health. However, in many cases the mechanisms underlying these beneficial effects are unknown, although a limited number of studies have suggested that bacteriocin production by *Bifidobacterium* may represent a key mechanism for preventing pathogen over-growth. Here, we used BAGEL4 and antiSMASH to identify putative bacteriocin sequences in the whole genome sequences of 33 *Bifidobacterium* strains isolated from infants participating in two clinical studies. We identified a novel non-lantibiotic bacteriocin from *Bifidobacterium longum* subsp. *infantis* LH_664, with 40% sequence homology to Lactococcin 972 from *Lactococcus lactis* subsp. *lactis*. The putative bacteriocin (Bifidococcin_664) was chemically synthesized and studied for antimicrobial and immune-modulatory activities. We determined it has discrete activity against *Clostridium perfringens* and it appears to have novel immune stimulatory activities, promoting macrophage phagocytosis and specific cytokine release. These data highlight strain-specific beneficial properties in the early life genus *Bifidobacterium*, and suggest avenues for development of novel and highly specific dual action antimicrobials, and possible probiotic strains, that are active against clinically important bacterial pathogens.

**Data summary:** Samples LH_9 to LH_666 were previously sequenced and deposited to ENA under accession numbers ERS2658025-ERS2658043. Samples LH_986 to LH_1052 are newly sequenced and deposited to NCBI under accession numbers SAMN24838598-SAMN24838611. Additionally, previously assembled publicly available sequences (n=7) were retrieved online from NCBI Genomes database. See Supplementary Table S1 for further details.

## Introduction

Antibiotics are critical for fighting serious bacterial infection, however their efficacy is declining due to increasing numbers of antibiotic-resistant bacteria (pathogens and commensals) (*1*). This global health crisis is further compounded by the lack of antibiotic development (*2*), thus there is a clear and urgent need for novel antimicrobials. Bacteriocins, which are ribosomally synthesized antimicrobial peptides produced by bacteria, are attractive alternatives to traditional antibiotics (3, 4). These naturally occurring compounds are active against other bacteria, and can be either narrow spectrum (i.e. targeting similar species) or broad spectrum (i.e. across diverse genera) (*5*), with the host bacterium resistant.

Bacteriocins are mainly categorized into three classes based on primary structure, molecular mass, thermal stability, and mode of action (*6, 7*). Class I undergo extensive post-translational modifications, which result in incorporation of non-standard amino acids (aa) (e.g. lanthionine and β-methyllanthionine) and are typically formed as 19 –50 aa chains. Nisin from *Lactococcus lactis* is a well-known example of a class I bacteriocin (*8*). Class II bacteriocins are unmodified peptides that can be divided into four subgroups: class IIa–d. Class IIa (or pediocin PA1-like) peptides are abundant, and comprised of 37-48 aa and positively charged (*9, 10*). They often exhibit strong activity against *Listeria* infections and are relatively heat stable. Heat labile bacteriocins make up the class III bacteriocins, however, there is limited information on these; *Escherichia coli* colicin and *Lactobacillus crispatus* helveticin M are a few known examples (*11*). Adherence of bacteriocins to the target microbe is mediated through docking molecules (e.g. Lipid II, phosphotransferase proteins IIC/D) (*11*), and their mode of action comprises either destabilisation of the proton motive force through pore formation, or inhibition of DNA-, RNA-, and protein-synthesis (*11*).

Previous genome mining studies have indicated that bacteriocin gene clusters are widespread within certain gut microbiota members (*12*). In early life, *Bifidobacterium* comprise the dominant microbiota genus in vaginally-delivered and breast-fed infants (*13*), with certain species more prevalent during this developmental window e.g. *Bifidobacterium longum* subsp. *infantis*. High levels of *Bifidobacterium* during infancy are associated with improved health, with certain species and strains important for the maintenance of the wider microbial ecosystem, prevention of enteric infections, and immune system programming (*13*). Numerous studies have indicated that disturbances in the early life bifidobacterial community are linked to an increased risk of serious infections (*14*), Very low levels or complete absence of gut-associated *Bifidobacterium* correlate with enhanced colonisation by potential pathogens such as *Enterococcus, Klebsiella*, and *Clostridium*, which can result in life-threatening conditions such as necrotising enterocolitis (NEC) and late onset sepsis (LOS) in preterm infants (*15*). Previous studies that have supplemented preterm infants with probiotics containing *Bifidobacterium* strains have highlighted that this protection may be linked to decreased gut pH, which is mediated by the production of short-chain fatty acids (SCFAs) after fermentation of breast milk-derived human milk oligosaccharides (HMOs), and/or via immune modulation (*16*). In addition, a limited number of studies have also suggested that bacteriocin production by *Bifidobacterium* may also represent a key mechanism for preventing pathogen over-growth (*17, 18*).

The production of bacteriocins by *Bifidobacterium* and their antimicrobial activity was first demonstrated in *Bifidobacterium bifidum* NCDC 1452. This bacteriocin, bifidin was shown to have activity against a range of Gram negative and Gram positive pathogens including *E. coli* and *Staphylococcus aureus* (*19*). Subsequently, bacteriocins bifilact Bb-12 and bifilong Bb-46, produced by *Bifidobacterium animalis* subsp. *lactis* Bb-12 and *B*. *longum* subsp. *longum* Bb-46 respectively, were reported to exhibit similarly strong activity against *S. aureus, E. coli, Salmonella enterica* serovar Typhimurium and *Bacillus cereus* (*20*). Although there are numerous reports on *Bifidobacterium*-associated bacteriocins, very few have been characterised in detail. Bifidocin B from *B. bifidum* NCFB 1454, bifidin I from *B. longum* subsp. *infantis* BCRC 14602 (partially sequenced), and the lantibiotic bisin from *B. longum* subsp. *longum* DJO10A, which share homology with their N-terminal region are the only ones characterised to date (*20*).

Many bacteriocin studies have focus on associated antimicrobial activity; however, these small peptides may also interact with host intestinal epithelial cells and cross the gut barrier to interact with underlying immune cells. Recent *in vitro* studies have suggested that Nisin A, plantaricin 423 (produced by *Lactobacillus plantarum* 423) and bacST4SA (produced by *Enterococcus mundtii* ST4SA) can traverse colonic Caco-2 monolayers (*21*), and at high concentrations Nisin A may activate neutrophils (*22*). Studies probing these potential immune-modulatory bacteriocin activities are very limited to those described above, and although extracellular proteins secreted by *Bifidobacterium* have been shown to stimulate host immune responses, there is currently no evidence suggesting that bacteriocins represent the active secretory component.

Although the studies described above provide important insights into *Bifidobacterium*-associated bacteriocins, further investigations are required to identify members of this beneficial early life genus for novel antimicrobials that may also have additional protective immune-modulatory activities. In the current study, we used BAGEL4 and antiSMASH to identify putative bacteriocin sequences in the whole genome sequences of 33 *Bifidobacterium* strains (*16, 23*). We identified a novel non-lantibiotic bacteriocin from *B. longum* subsp. *infantis* LH_664 (Bifidococcin_664), which when chemically synthesised was found to have discrete activity against *C. perfringens* and immune modulatory activities.

## Results

### General features of analysed genomes

Genomic features of strains LH_9 to LH_666 have previously been described (*23*). The genome sizes of the newly sequenced isolates (LH_986 to LH_1052) ranged from 1.94 Mb for LH_990 to 2.4 Mb for LH_1004, which corresponded to 1,627 and 2,117 ORFs, respectively (**Supplementary Table 1**). The G+C% content ranged from 58.59% for LH_1052 to 62.76% for both LH_1016 and LH_1036, consistent with previously published data (*24*).

Phylogenomic position and genetic relatedness of the isolates included in this study to the recognised human-associated *Bifidobacterium* species were determined based on the average nucleotide identity (ANI) values and the alignment-free whole genome analysis (**Fig. 1**). The results indicated the clustering of isolates into five phylogenetic groups, namely *Bifidobacterium pseudocatenulatum, B. animalis* subsp. *lactis, B. bifidum, Bifidobacterium breve*, and *B. longum*, with newly sequenced isolates identified as members the *B. bifidum, B. breve* and *B. longum* taxa.

**Figure 1:**
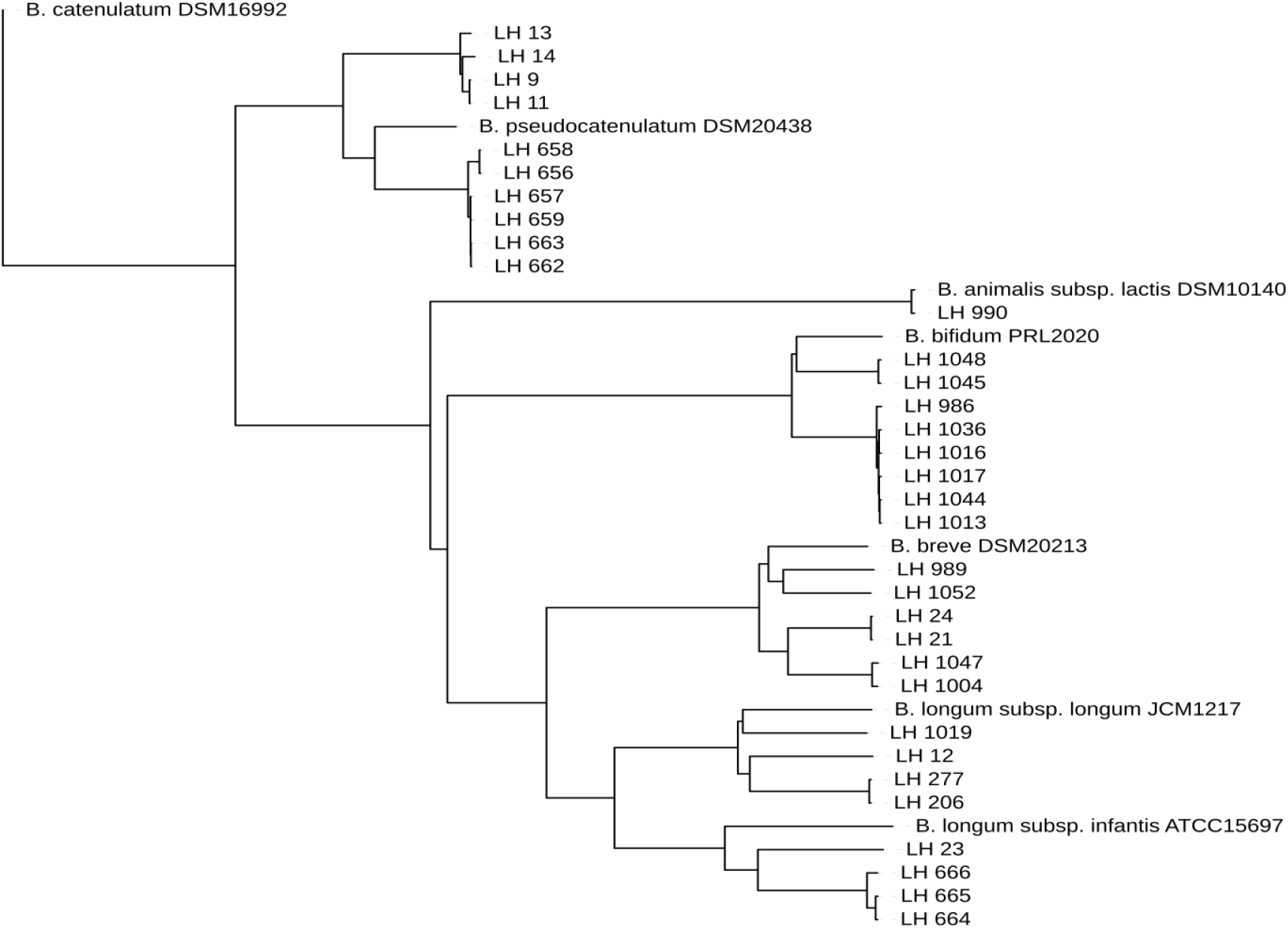
Phylogenomic position and genetic relatedness of isolates used in this study. Alignment-free composition approach (k=6) indicated the clustering of isolates into five phylogenetic groups, namely *Bifidobacterium pseudocatenulatum, Bifidobacterium animalis* subsp. *lactis, Bifidobacterium bifidum, Bifidobacterium breve*, and *Bifidobacterium longum*.

### Genome mining for putative bacteriocin synthesizing genes

Thirty-three bifidobacterial genomes were analysed for the presence of putative bacteriocin synthesizing genes using antiSMASH 5 and BAGEL4, which run based on Hidden Markov Models (HMM) to detect broad array of secondary metabolites from bacterial genomes (*25*). This analysis revealed the presence of bacteriocin encoding genes in five bifidobacterial isolates. Putative bacteriocins identified in *B. longum* subsp. *longum* LH_1019 and *B. longum* subsp. *infantis* LH_23 showed high homology with known bacteriocins mercasidin and michiganin-A, respectively. The other three strains, namely *B. longum* subsp. *infantis* LH_664, *B. longum* subsp. *longum* LH_665, and *B. breve* LH_666, were found to putatively encode sactipeptide (Class I bacteriocin), however the predicted genes did not show high similarities with known bacteriocins (**Table 1**). In addition, the nucleotide blast alignment of the predicted structural genes also did not indicate homology to GenBank databases (represented as hypothetical protein). Therefore, these predictions were considered particularly interesting, and we hypothesised that they may encode novel bacteriocins. Sequence re-analysis using antiSMASH revealed the presence of putative bacteriocin synthesis genes in these strains (**Fig. 2A**), with same sequence matching to bacteriocin Lactococcin_972 (belonging to the Lactococcin972 family). In-depth analysis using Pfam and InterProScan searches confirmed that *B. longum* subsp. *infantis* LH_664 and *B. breve* LH_666 each encode a 134-aa peptide, while *B. longum* subsp. *longum* LH_665 encodes a 139-aa peptide, with 34-aa leader sequence and over 100-aa bacteriocin peptide (**Fig. 2B**). Further alignment analysis of the 3 strains showed that 63 residues (47% of the sequence) were homologous to the known bacteriocin Lactoccocin_972 produced by *L. lactis* subsp. *lactis* and the results of the molecular modelling with phre^2^ also confirmed 99.7% identity of those residues to the single highest scoring template, Lactoccocin_972 (**Fig. 2C**). Therefore, the putative bacteriocins were named Bifidococcins, tagging the respective strain number at the end. Although the three bacteriocins have different leader sequences, the Lactococcin_972 sequences encoded within the complete cluster may have been acquired through horizontal gene transfer from a *Lactococccus* species. As the three strains and encoded bacteriocins showed the same structural properties *in silico*, we opted to select one for subsequent analysis and characterisation. We decided to focus on the bacteriocin encoded in *B. longum* subsp. *infantis* LH_664, given this *Bifidobacterium* subspecies is known to play a key role in the early life gut microbiota and microbe-microbe interactions (*26, 27*). The complete, 134-aa Bifidococcin_664 was chemically synthesised through solid phase synthesis and obtained as > 95% pure compound, checked through high performance liquid chromatography (Lifetein, LLC, USA). Bifidococcin_664 is predicted to be cationic (using compute pI/Mw tool), with a putative net charge of 2.2 (theoretical pI = 8.94) and a molecular weight of 14972.96 g/mol (14%). We determined that the solubility range is 3mg/ml with gentle heating at 65°C for 5 min in water. This peptide also had high solubility in Dimethyl sulfoxide (DMSO), and low pH (3.5), despite this we used water to ensure compatibility with downstream *in vitro* tests (data not shown).

**Table 1.** Results of bacteriocin screens performed with antiSMASH and BAGEL4 on the 33 bifidobacterial genomes.

| S.no | *Bifidobacterium* species identified | Isolation ID | Source (Preterm = <37 weeks gestation) | Dietary source | Bacteriocin identified (antiSMASH/BAGEL4) |
| --- | --- | --- | --- | --- | --- |
| 1 | *Bifidobacterium bifidum* | LH_986 | Preterm infant gut | Breast milk | No |
| 2 | *Bifidobacterium bifidum* | LH_1017 | Preterm infant gut | Breast milk | No |
| 3 | *Bifidobacterium bifidum* | LH_1045 | Preterm infant gut | Breast milk | No |
| 4 | *Bifidobacterium bifidum* | LH_1048 | Preterm infant gut | Breast milk | No |
| 5 | *Bifidobacterium bifidum* | LH_1044 | Preterm infant gut | Breast milk | No |
| 6 | *Bifidobacterium bifidum* | LH_1036 | Preterm infant gut | Breast milk | No |
| 7 | *Bifidobacterium bifidum* | LH_1013 | Preterm infant gut | Breast milk | No |
| 8 | *Bifidobacterium bifidum* | LH_1016 | Preterm infant gut | Formula fed containing HMOs | No |
| 9 | *Bifidobacterium breve* | LH_989 | Preterm infant gut | Breast milk | No |
| 10 | *Bifidobacterium breve* | LH_990 | Preterm infant gut | Breast Milk | No |
| 11 | *Bifidobacterium breve* | LH_1004 | Preterm infant gut | Formula fed containing HMOs | No |
| 12 | *Bifidobacterium breve* | LH_1047 | Preterm infant gut | Breast milk | No |
| 13 | *Bifidobacterium breve* | LH_1052 | Preterm infant gut | Breast milk | No |
| 14 | *Bifidobacterium longum* subsp. *longum* | LH_1019 | Preterm infant gut | Breast milk | Mercasidin |
| 15 | *Bifidobacterium pseudocatenulatum* | LH_9 | Full term infant gut | Breast milk | No |
| 16 | *Bifidobacterium pseudocatenulatum* | LH_11 | Full term infant gut | Breast milk | No |
| 17 | *Bifidobacterium longum* subsp. *longum* | LH_12 | Full term infant gut | Breast milk | No |
| 18 | *Bifidobacterium pseudocatenulatum* | LH_13 | Full term infant gut | Breast milk | No |
| 19 | *Bifidobacterium pseudocatenulatum* | LH_14 | Full term infant gut | Breast milk | No |
| 20 | *Bifidobacterium breve* | LH_21 | Full term infant gut | Breast milk | No |
| 21 | *Bifidobacterium longum* subsp. *infantis* | LH_23 | Full term infant gut | Breast milk | Michiganin-A |
| 22 | *Bifidobacterium breve* strain NRBB56 | LH_24 | Full term infant gut | Breast milk | No |
| 23 | *Bifidobacterium longum* | LH_206 | Full term infant gut | Breast milk | No |
| 24 | *Bifidobacterium longum* subsp. *longum* | LH_277 | Full term infant gut | Breast milk | No |
| 25 | *Bifidobacterium pseudocatenulatum* | LH_656 | Full term infant gut | Breast milk | No |
| 26 | *Bifidobacterium pseudocatenulatum* | LH_657 | Full term infant gut | Breast milk | No |
| 27 | *Bifidobacterium pseudocatenulatum* | LH_658 | Full term infant gut | Breast milk | No |
| 28 | *Bifidobacterium pseudocatenulatum* | LH_659 | Full term infant gut | Breast milk | No |
| 29 | *Bifidobacterium pseudocatenulatum* | LH_662 | Full term infant gut | Breast milk | No |
| 30 | *Bifidobacterium pseudocatenulatum* | LH_663 | Full term infant gut | Breast milk | No |
| 31 | *Bifidobacterium longum* subsp. *infantis* | LH_664 | Full term infant gut | Breast milk | sactipeptides |
| 32 | *Bifidobacterium longum* subsp. *longum* GT15 | LH_665 | Full term infant gut | Breast milk | sactipeptides |
| 33 | *Bifidobacterium breve* | LH_666 | Full term infant gut | Breast milk | sactipeptides |

**Figure 2:**
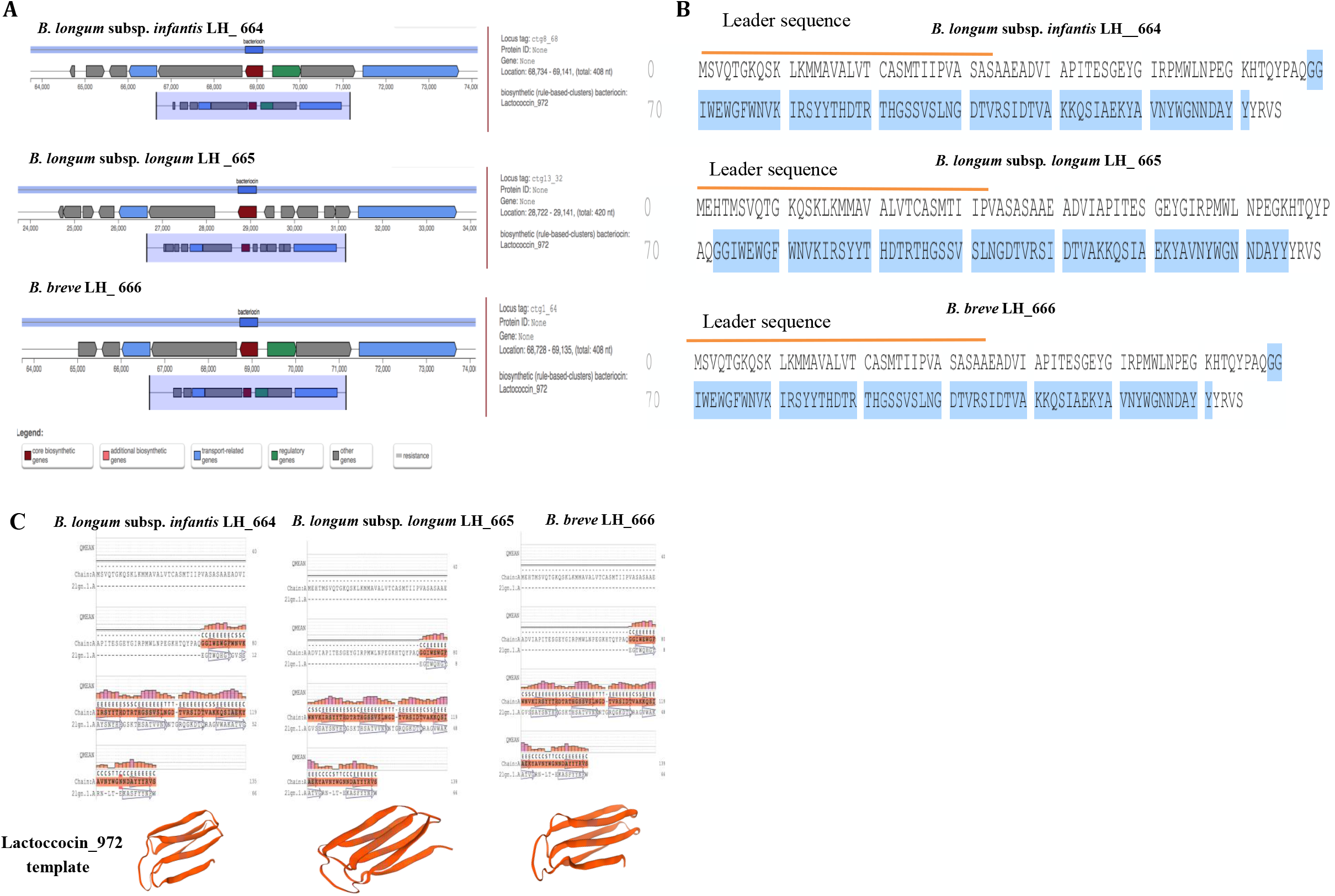
*In silico* bacteriocin analysis. **(A)** Results of antiSMASH analysis showing bacteriocin biosynthetic gene cluster (putative and functional genes) in the genomes of *B. longum* subsp. *infantis* LH_664, *B. longum* subsp. *longum* LH_665, *B. breve* LH_666 and similarity matches with known bacteriocin Lactococcin_972. **(B)** Results of Pfam and InterProScan sequence analysis showing all three predicted bacteriocins have a leader sequence (34-aa) (marked by orange line) and amino acid sequences (64-aa) (highlighted in blue) homologous to those of Lactococcin_972. **(C)** Results of molecular modeling with phre^2^ confirming that 64-aa regions of three predicted bacteriocins show 99.7% identity to the single highest scoring template Lactoccocin_972, with the structure of a β-sandwich comprising two three-stranded antiparallel β-sheets displayed underneath alignment.

### Impact of Bifidococcin_664 on microbial growth

In order to determine any potential antimicrobial properties of Bifidococcin_664, we selected two important species, generally associated with preterm NEC, *Enterococcus faecium* and *Clostridium perfringens*, alongside strains expected to be resistant to Lactococcin 972: *Lactococcus garvieae* and *Lactobacillus animalis* (*28, 38*). Initial studies with concentrations below 100 μM did not result in growth impairment of any of the tested bacterial strains (data not shown). Incubations with doses above 100 μM (at 133μM) also did not negatively affect growth of *L. garvieae* LH_269, *L. animalis* LH_283, and *E. faecium* LH_302 (**Fig. 3A**). However, we did observe that *C. perfringens* LH_19 showed a modest reduction in exponential growth when Biffidococcin_664 was present at 100uM suggesting this bacterium may be sensitive (**Fig. 3A**). Further, growth assays with increasing concentrations of Bifidococcin_664 (100, 200 and 300 μM), indicated a gradual decrease in the exponential growth phase (from doubling time 40 minutes) (**Fig. 3B and Fig. 3C**). Notably, at 300μM we observed an extended lag phase, with a doubling time of approximately 8.7 hours (**Fig. 3B and Fig. 3C**). These data indicate that Bifidococcin_664 specifically inhibits the pathogen *C. perfringens* LH_19 across a range of concentrations.

**Figure 3.**
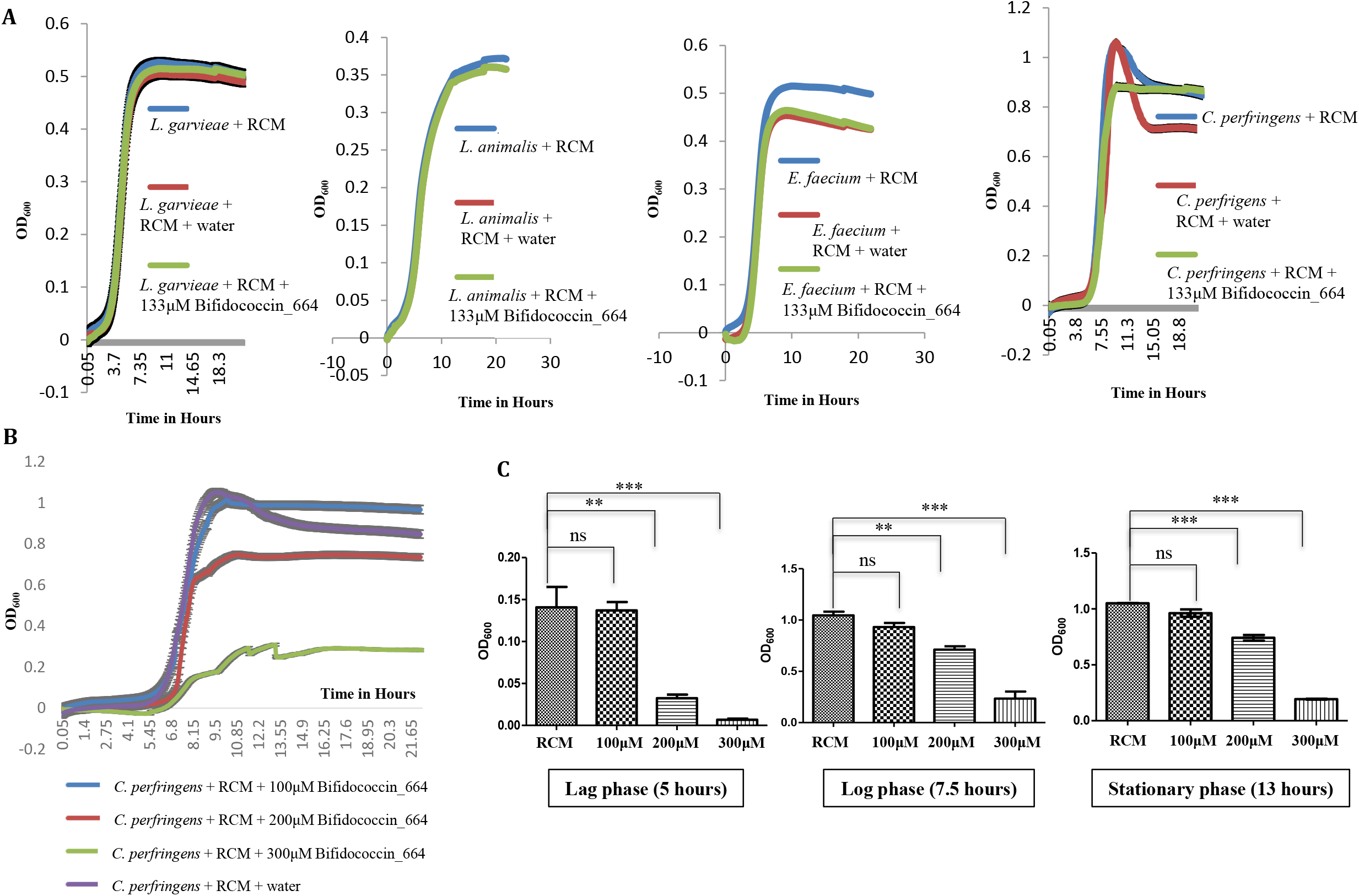
Growth studies of Bifidococcin_664 antimicrobial activity against different bacterial strains. **(A)** The initial tested concentration (130 μM) did not impact growth (OD_600_) of *L. garvieae* LH_269, *L. animalis* LH_283, and *E. faecium* LH_302, but did have modest antimicrobial activity against *C. perfringens* LH_19. **(B)** Further OD_600_ growth curves of *C. perfringens* LH_19 treated with increasing concentrations of Bifidococcin_664 (i.e. 100, 200 and 300 μM) resulted in a gradual decrease during exponential growth. **(C)** Graph showing reduction of growth (representative OD_600_) at different growth phases of *C. perfringens* LH_19 upon treatment with different concentrations of Bifidococcin_664 and control (RCM) (mean ±SD), p values ns=>0.05, <0.0001, 0.0001.

### Determination of Bifidococcin_664 immune modulatory properties

Previous work has suggested that bacteriocins may also influence immune responses. Therefore, to test if Bifidococcin_664 has dual activity, both antimicrobial and immune modulatory, we initially used a macrophage reporter cell line that displays colorimetric activity in response to NF-κB stimulation (i.e. THP1-Blue™ cells). Cells were stimulated for 24 hours with increasing doses of Bifidococcin_664 (17.368 μM, 43.4 μM and 86.4 μM) along with LPS and water as positive and negative control respectively, with activation of NF-κB measured using QUANTI-Blue™. A concentration-dependent fold increase (2, 3.6 and 6) of NF-κB activation was observed in Bifidococcin_664 treated THP1-Blue cells, compared to the negative control (**Fig. 4A** and **Fig. 4B**).

**Figure 4.**
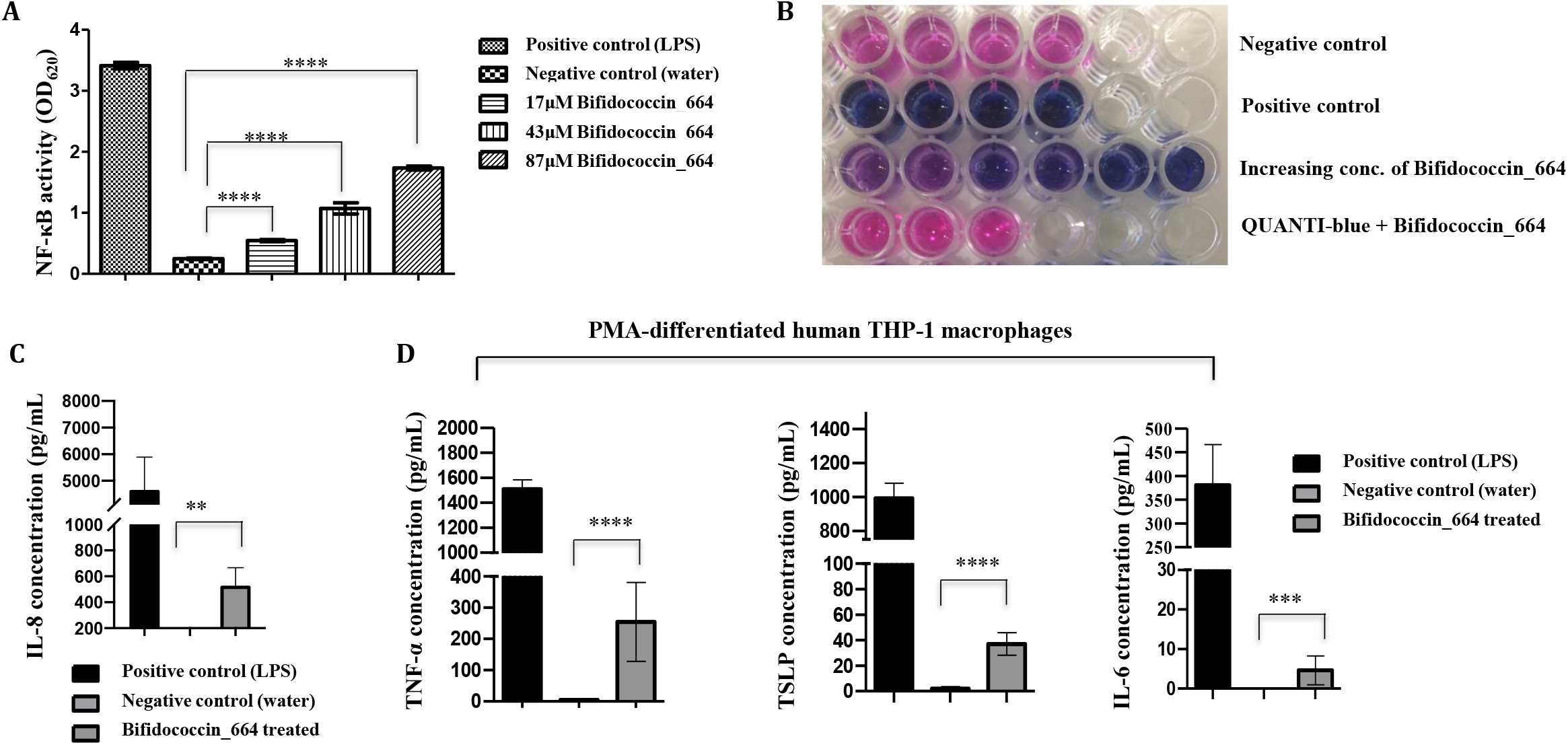
Impact of increasing concentrations of Bifidococcin_664 on macrophage-like cell line. (THP1-Blue cells at 24 hours). **(A)** The level of NF-κB activation was measured and **(B)** quantified using QUANTI-Blue (see materials and methods for further details). Data is from 3 independent experiments (mean ±SD) and statistical significance (ANOVA) calculated (minus the negative control), with P value is <0.0001. **(C)** Level of IL-8 released from Bifidococcin_664 treated THP-1 blue monocytes (P <0.003). Levels of cytokines detected from Bifidococcin_664 treated PMA-differentiated human THP-1 macrophages **(D)** TNF-α (P <0.001), TSLP (P <0/001), and IL-6 (P <0.002). Data is from two independent experiments (mean ±SD), with statistical significance (ANOVA) calculated over the negative control.

To further determine potential immune modulatory abilities of Bifidococcin_664, THP1-Blue monocytes, alongside PMA-differentiated human THP-1 macrophages and human colon carcinoma Caco-2 cells were assayed for cytokine production (IL-6, IL-15, IL-8, TSLP and TNF-α). THP-1 blue monocytes released increased levels of IL-8 (400 fold vs. negative control) after Bifidococcin_664 treatment, while secretion of other cytokines (IL-6, IL-15, TSLP and TNF-α) was not detectable (**Fig. 4C**). PMA-differentiated human THP-1 macrophages produced significant levels of pro-inflammatory cytokines TNF-α and TSLP (70 and 30 fold higher vs. negative control) (**Fig. 4D**), along with increased (albeit low) concentrations of IL-6 in response to Bifidococcin_664 treatment (**Fig. 4D**) (all p values <0.05). However, there were no traces of IL-15. Interestingly, we did not observe any impact of Bifidococcin_664 treatment on CaCo2 cells (data not shown), suggesting the mode of action may be restricted to professional immune cells i.e. macrophages, rather than intestinal epithelial cells.

### Phagocytosis of *S*. Typhimurium in Bifidococcin_664 treated PMA-differentiated human THP-1 macrophages

The activation and inflammatory status of immune cells correlates with their anti-pathogen activity, therefore we also determined if Bifidococcin_664 was able to enhance phagocytosis in PMA-differentiated human THP-1 macrophages pre-treated for 15 hours. We used a gentamycin survival assay with *S*. Typhimurium 3580 (which was resistant to direct antimicrobial activities of Bifidococcin_664 (**Supplementary Fig. 1**). We observed enhanced phagocytosis of *S*. Typhimurium 3580 in Bifidococcin_664 pre-treated macrophages, when compared to non-treated macrophages (22-fold higher) 1 hour post incubation (**Fig. 5**). However, a longer incubation (2.5 hours) did not enhance this activity, as we observed the expected killing of *S*. Typhimurium across all experimental groups (**Fig. 5**). These data indicate that Bifidococcin_664 may enhance the rapid internalisation of *Salmonella*, but it does not influence the ability of macrophage to efficiently lyse/kill internalised bacteria.

**Figure 5:**
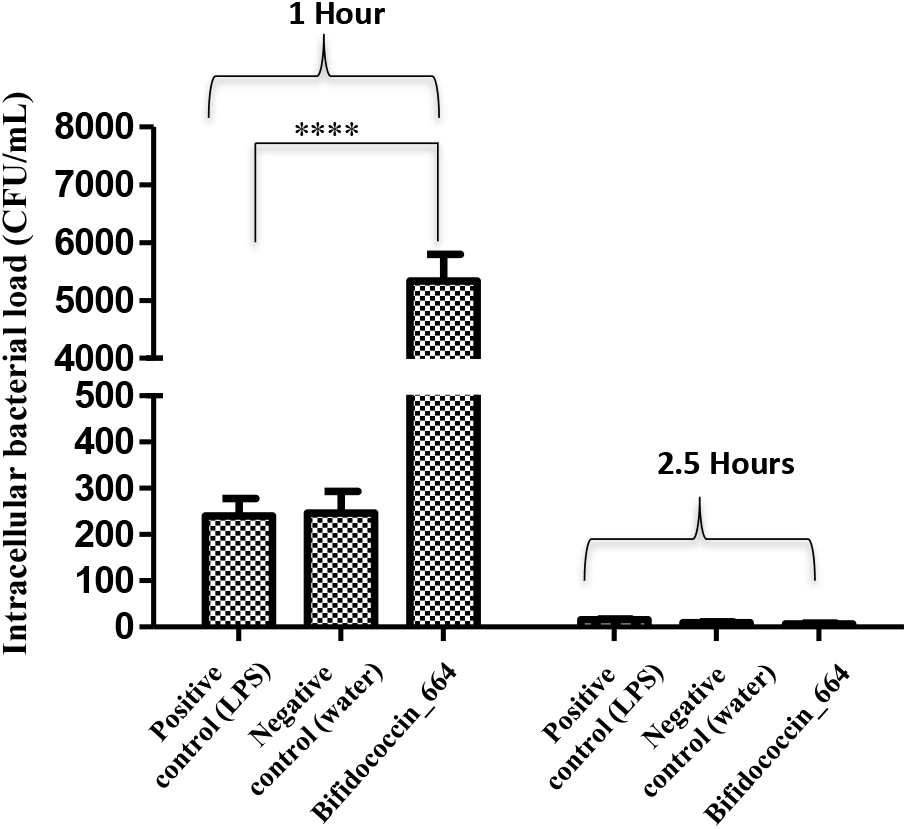
Survival assay of *S*. Typhimurium 3580 in Bifidococcin_664 treated PMA-differentiated human THP-1 macrophages. Enhanced phagocytosis and survival of *S. Typhimurium* 3580 was observed after 1 hour of incubation. At longer incubation (2.5 hours) the intracellular level of *S*. Typhimurium 3580 was reduced, and Bifidococcin_664 treated PMA-differentiated human THP-1 macrophages did not appear to further exhibit killing activity. Data are means ± SD from three independent experiments. Statistical significance was determined using two way ANOVA and the p value is <0.0001

## Discussion

Within the gut, bacteriocin-producing bacteria provide an important benefit for the host via colonisation resistance mechanisms. These anti-microbial peptides may inhibit over-growth of microbiota members that can act as opportunistic pathogens under certain conditions and/or bacteriocins may prevent colonisation of invading pathogenic bacteria. This is particularly important in at-risk groups such as neonates (including preterm infants) who do not have a mature immune system and lack a robust early life gut microbiota, which limits their ability to mount effective anti-pathogen responses. Moreover, these infants are often exposed to multidrug resistant bacteria, and they harbour a high number of commensal bacteria that encode AMR genes (*29, 30*), thus identification and development of novel anti-microbial strategies, like bacteriocins, are crucial. However, to date, our understanding and characterisation of bacteriocin producing microbiota members and bacteriocin function is somewhat limited.

In this study, we focused on profiling different species and isolates of the key early life genus *Bifidobacterium*, mining 33 genomes for the presence of bacteriocin synthesising genes. Our initial *in-silico* screen with BAGEL4 identified two known functional lantibiotics, and three unknown sactipeptides. A functional bacteriocin includes a structural gene encoding a precursor, genes encoding an immunity protein, an ATP-binding cassette transporter, and a protein for extracellular translocation (*31*). Based on our initial analysis we did not find these genes surrounding the identified sactipeptides, thus we regarded them as potential false positives. However, subsequent ‘re-mining’ of the genomes with antiSMASH 5 predicted putative genes required for synthesis of a functional bacteriocin in the genomes of bifidobacterial isolates LH_664, LH_665, and LH_666. Based on this prediction, and the 47% sequence homology with known bacteriocin Lactococcin_972, we named all three Bifidococcins (with isolate IDs as peptide descriptor) (**Fig. 2**). Although all predicted Bifidococcins showed the same identity values to a known bacteriocin Lactococcin_972 (99.7% homology with 63 residues), further *in silico* analysis indicated isolate-specific differenced in peptide sequences (a further 134 residues in Bifidococcin_664, a different predicted molecular weight (14.9 kDa), a leader sequence of 34 amino acids, MSVQTGK as N-terminus and AY YYRVS as C-terminus sequences, in addition to the 60% unmatched sequence homology with any of the known bacteriocins. Therefore, based on these data (and downstream characterisation work) we propose that Bifidococcin is a novel bacteriocin belonging to the *Bifidobacterium* genera. Importantly, our *in silico* analysis also highlighted key isolate-level differences with regard to the presence of putative bacteriocin synthesis genes, as the majority of the screened isolates were not predicted to encode any putative antimicrobial peptides. Thus, where anti-pathogen responses are required, genome-wide pre-screening to identify the most ‘promising’ isolates should be considered for the rational selection of probiotic use in the clinic.

Bacteriocins are known to have a range of antimicrobial activities, which may be beneficial for microbiota members for efficient colonisation and ecosystem structuring during the early life developmental window (*26, 32*). These compounds also provide a key protective effect for the host, potentially limiting over-growth of pathobionts in the gut (*4, 33, 34*). Through solid phase synthesis, we synthesised the bacteriocin of *B. longum* subsp. *infantis* LH_664 (Bifidococcin_664) and tested its ability to inhibit bacterial growth. Although its sequence showed 47% homology to Lactococcin_972, we observed that tested *Lactococcus* and *Lactobacillus* isolates were resistant (i.e. displayed no growth defects) to Bifidococcin_664. Indeed, host isolates or similar genetic variants are often resistant to their ‘native’ bacteriocin, and thus lysis (*35*). We also explored the ability of Bifidococcin_664 to inhibit growth of key preterm pathogens that are associated with severe and often deadly gastrointestinal diseases such as NEC. Preterm infants have a radically altered gut microbiota, often lacking *Bifidobacterium* members, but enriched in potential pathogens such as *Enterococcus* and *Clostridium*, which are associated with higher risk of NEC (*36, 37*). Notably, probiotic supplementation (with bifidobacterial strains) is often used to ‘restore’ the early life gut microbiota, and decrease NEC incidence (*16, 38, 39*). Although we did not observe any activity against *E. faecium*, we did observe inhibition of *C. perfringens* LH_19, which was dose dependent (**Fig. 3**), suggesting potential narrow spectrum activity of Bifidococcin_664 towards this NEC-associated species (*38*). Interestingly, our recent studies in a preterm cohort have indicated supplementation with *Bifidobacterium* is associated with concurrent reduction in *Clostridium*, with this probiotic treatment also being associated with a more than 50% reduction in the incidence of NEC (and of late-onset sepsis), which may link to additional benefits afforded though bacteriocin production in the preterm gut environment (*16*). Further screening assays against numerous microbiota members and pathogens would be required to determine the full antimicrobial activity parameters of Bifidococcin_664 in future studies. It is also unclear from our initial studies what the mode of action of this bacteriocin is. Therefore, additional studies would be required to characterise this and determine if the antimicrobial activity is similar to Lactococcin 972, which acts through inhibition of septum formation resulting in structural cellular changes that eventually lead to lysis and cell death (*28*).

Previous work has indicated that bacteriocins are non-toxic to eukaryotic cells, and limited number of reports suggested that these peptides might modulate host cell responses (*4, 21, 40*). Although *Bifidobacterium* is abundant in the colon, we did not observe any modulation of Caco2 responses following treatment with Bifidococcin_664, suggesting these host cells may not play a role in bacteriocin-host crosstalk (albeit these are a colonic cancer cell line and thus are not fully representative of the colonic epithelium). Surprisingly, Bifidococcin_664 induced the central immune mediator NF-κB in THP1 monocytes, in tandem with IL-8 release, suggesting this bacteriocin may also have immune-modulatory activity. Although monocytes (and other immune cells) are found within the underlying mucosa of the gut, previous studies have indicated that bacteriocins can transverse the epithelial barrier and thus could be expected to interact with various immune cells (*21*). Moreover, to further assess the immunomodulatory properties of Bifidococcin_664, we additionally utilised PMA-differentiated human THP-1 macrophages, which also responded to treatment as indicated by differential cytokine release (IL-6, IL-8, TSLP, and TNF-α). This pro-inflammatory milieu may also correlate with the enhanced internalisation of *Salmonella* as observed in the phagocytosis assay, via ‘priming’ of macrophage anti-infection responses (**Fig. 5**). Indeed, this rapid uptake (observed at 1-hour post inoculation) may serve to limit over-growth of rapidly replicating and invasive bacterial pathogens in the gut. Further investigations using *in vivo* infection models could be used to determine the role of these responses in detail, with additional studies exploring potential peptide-receptor interactions required to elucidate the signalling pathways involved.

## Conclusions

To our knowledge, this is the first report of a dual natured bacteriocin from the early life microbiota member *Bifidobacterium (B. longum* subsp. *infantis* LH_664). Bacteriocins have several advantages over traditional antibiotics for infection treatment as they are more specific in their killing, and avoid the collateral damage often associated with antibiotic use across the wider gut microbiota (*33*), which in itself is associated with higher infection risk. Moreover, bacteriocins in combination with antibiotics have further advantages as they reduce the concentration of antibiotics needed (*33*), particularly against those bacteria residing in biofilms (*41*). Delivery of whole bacteria to the gut and targeted production of bacteriocins may be problematic, thus synthesis of discrete peptide(s) may provide a treatment alternative, including the ability to administer these molecules to combat systemic infections, whilst a gut-derived antimicrobial may have a less potent effect. The fact that our initial studies also suggest this bacteriocin may augment anti-infection responses in professional immune populations (i.e. macrophages) also suggests the utility of Bifidococcin_664 for hard to treat (i.e. multidrug resistant) and invasive bacterial pathogens. Further in-depth studies focused on the *in vitro* and *in vivo* characterisation of this novel bacteriocin may allow development of novel antimicrobial therapies for treatment of serious neonatal infections.

## Author contributions

SGJ and LJH designed the overall project and experiments. SGJ carried out the majority of the studies, analysed the data, and drafted the manuscript. MK performed the phylogenetic analysis. DP performed the cytokine panel assay. AG and NB completed and supervised the CaCo2 cell assays. AJ contributed to the ELISA assays. MAEL, IJO’N, RK and CAG isolated the strains used in this study. PC provided clinical oversight for the original faecal samples used for *Bifidobacterium* isolation. LJH acquired funding, supervised the project, contributed to data analysis, and drafted the manuscript. All authors provided feedback on the manuscript draft prior to publication.

## Acknowledgments

This research was supported in part by the NBI Computing infrastructure for Science (CiS) group through the provision of a High-Performance Computing (HPC) Cluster. L.J.H. is supported by Wellcome Trust Investigator Awards 100974/C/13/Z and 220876/Z/20/Z; the Biotechnology and Biological Sciences Research Council (BBSRC), Institute Strategic Programme Gut Microbes and Health BB/R012490/1, and its constituent projects BBS/E/F/000PR10353 and BBS/E/F/000PR10356, a BBSRC Norwich Research Park and a Bioscience Doctoral Training grant no. BB/M011216/1 (supervisor LJH, student MK), and a previous Crohn’s and Colitis UK award [M/15/6]. We thank the sequencing team at both Wellcome Trust Sanger Institute and Quadram Institute Bioscience for genome sequencing.

## Materials and Methods

### Bacterial isolation and strains

Infant faeces were collected from freshly soiled nappies and plated on Reinforced Clostridial Medium (RCM) (Oxoid™, Thermo Fischer Scientific, UK) supplemented with 0.05 mg/mL mupirocin and 0.05 mg/mL of L-cysteine (both from Sigma-Aldrich, UK) within four hours of collection. All *Bifidobacterium* isolates were streaked for purity and all strains were grown at 37°C in RCM and de Man Rogosa and Sharpe (MRS) (BD Difco™, BD Biosciences, USA) media in an anaerobic chamber (Whitley DG250, Don Whitley Scientific, UK), unless otherwise specified. All samples were collected according to the Quadram Institute Bioscience Ethics Committee, and sample collection was in accordance with protocols laid out by the National Research Ethics Service (NRES) approved UEA/QIB Biorepository (Licence no: 11208).

### Genomic DNA extraction

Overnight bacterial cultures (10mL) were grown and used for phenol-chloroform DNA extraction. Briefly, bacterial pellets were re-suspended in 2□ml of 25% sucrose in 10□mM Tris and 1□mM EDTA at pH 8.0. Cells were then treated using 50□μl of 100□mg/ml lysozyme (Roche Molecular Systems). Furthermore, 100□μl of 20□mg/ml Proteinase K (Roche Molecular Systems), 30□μl of 10□mg/ml RNase A (Roche Molecular Systems), 400□μl of 0.5□M EDTA (pH 8.0) and 250□μl of 10% Sarkosyl NL30 (Sigma-Aldrich) were added to the lysed bacterial suspension. The samples were then incubated on ice for 2 hours, and then placed in a 50°C water bath overnight, followed by three rounds of Phenol:Chloroform:Isoamyl Alcohol (25:24:1) (Sigma-Aldrich) extraction using Qiagen MaXtract High Density tubes (Qiagen), and two rounds of extractions with Chloroform:Isoamyl Alcohol (24:1) (Sigma-Aldrich) to remove residual phenol before performing ethanol precipitation and wash (in 70% ethanol). Once dry, genomic DNA pellets were resuspended in 300□μl of 10□mM Tris (pH 8.0) and quantified using Qubit dsDNA BR assay kit.

### Whole genome sequencing

Samples LH_9 to LH_666 were previously sequenced and deposited to ENA under accession numbers ERS2658025-ERS2658043 (*23*). Samples LH_986 to LH_1052 were sequenced on llumina MiSeq with read length 2 × 150bp (paired-end reads, with average sequencing coverage of 64x). Draft genome assemblies were generated using SPAdes v3.11.1 (*42*). Samples LH_986 to LH_1052 are deposited to NCBI under accession numbers SAMN24838598-SAMN24838611. Additionally, previously assembled publicly available sequences (n=7) were retrieved online from NCBI Genomes database (*43*) (Supplementary Table S1).

### Alignment-free whole genome phylogeny

Genomes of 33 *Bifidobacterium* strains were subjected to Prokka v1.14 (*44*) to obtain annotated FASTA files. Alignment-free whole genome phylogeny was reconstructed using CVTree v5.0, with parameters k = 6 on amino acid sequences (*45*). Interactive Tree of Life was used to visualise phylogeny (*46*) (http://itol.embl.de/).

### Average nucleotide identity (ANI)

Python 3 module pyANI using BLASTN+ was employed to calculate the average nucleotide identity between genomes (*47*). Identity□>95% was used as cut-off for species delineation.

### Bacterial culturing conditions

The primary culture of *B. longum* subsp. *infantis* LH_664 was inoculated in 5 ml of RCM supplemented with L-cysteine (0.05 mg/mL) and incubated for 24 hours at 37°C in an anaerobic chamber. Pathogens and commensals (in-house clinical isolates) to be tested for antimicrobial assays, namely *Lactococcus garvieae* LH_269 *Lactobacillus animalis* LH_283, *Enterococcus faecium* LH_302, *Clostridium perfringens* LH_19, were maintained in RCM in anaerobic conditions at temperature 37°C.

### *In silico* identification and visualization of bifidococcin664

BAGEL4 (*48*) and antiSMASH v5. (*49*) were used to screen the twelve *Bifidobacterium* genomes for putative bacteriocin biosynthesis gene clusters. Additional analyses of the predicted bacteriocin genes were carried out using BLASTP, interpro scan and Pfam searches (*50*) (*51*). Bacteriocin structural characteristics were visualized using Phre^2^, RCSB protein data bank, and molecular weight and isoelectric point calculated using Compute pI/Mw tool (UniProt Knowledge base).

### Chemical synthesis of bifidococcin664

The predicted bacteriocin of *B. longum* subsp. *infantis* LH_664 with following sequence of amino acids MSVQTGKQSKLKMMAVALVTCASMTIIPVASASAAEADVIAPITESGEYGIRPMWLNPEGKHTQYPAQGGIWEWGFWNVKIRSYYTHDTRTHGSSVSLNGDTVRSIDTVAKKQSIAEKYAVNYWGNNDAYYYRVS was chemically synthesized by LifeTein (South Plainfield, NJ, USA).

### Growth kinetic assay in the presence of the bifidococcin664

To assess the inhibitory effect of Bifidococcin_664 growth kinetics were performed. An experimental outline was adapted as described, with some modification(*52*). Briefly, 10 μl of test strain suspensions (OD_600 nm_ = 0.4) was inoculated into 280 μl (5% level (v/v) inoculum) of each treatment combination (culture medium and bacteriocin/culture medium and control) in NUNC 96 multi-well plates. The volume of bacteriocin and buffer control added to the wells was 10 μl respectively. Growth was determined using microplate spectrophotometer (Cerillo, USA) placed in anaerobic chamber that was set to 37°C. Assays were performed three times, and data averaged.

### THP-1 NF-kB reporter cell line and reporter assay

THP1-Blue NF-κB cells (Invivogen, UK) were revived in growth medium RPMI 1640 with 2 mM L-glutamine, 25 mM HEPES (Merck, UK), 10% heat-inactivated foetal bovine serum (Gibco, UK), 100 μg/ml Normocin (Invivogen, UK), and Pen-Strep (100 U/ml-100 μg/ml) (Gibco, UK). These cells have a construct containing a secreted embryonic alkaline phosphatase (SEAP) gene induced by the nuclear factor κ-light-chain-enhancer of activated B cells (NF-κB) transcription factor. Like other monocytes, THP1-Blue cells, respond to ligands, such as peptides, endotoxins, and pathogenic stimulus via toll-like receptors (TLR). Upon TLR stimulation, NF-κB is activated; subsequently, the reporter enzyme is secreted into the cell culture medium, which is then quantified using the colorimetric reagent QUANTI-Blue™.

The Quanti-blue assay was conducted as per manufactures instructions (Quanti-Blue, Invivogen, UK). In brief, once 70% confluence was reached, Blasticidin (Invivogen, UK) treated THP1-Blue NF-κB cells were transferred from cell culture flasks and seeded at 1×10^5^ per well in a 96-well tissue culture plate. Cells were then stimulated with 17.368 μM, 43.402 μM, and 86.8403 μM of bifidococcin664 separately and incubated for 18 hours in a humidified incubator at 37° C with 5% CO_2_. LPS provided by manufacturer was used as a positive control. 20 μl aliquots of cell culture medium were removed after incubation and added to plates containing 180 μl of pre-warmed Quanti-Blue detection reagent per well as per manufacturer’s instructions. Colour was allowed to develop for 1 hour, and absorbance was read at 650 nm in microplate reader (Biotec, USA) (Ref).

### Macrophage assay

*Salmonella enterica* serovar Typhimurium was grown in Brain Heart Infusion (BHI) (Oxoid, UK) broth + 1.5% agar and incubated overnight at 37°C in an anaerobic incubator. THP-1 monocytes were obtained from ATCC (TIB-202) and maintained in RPMI (Gibco) plus 10% heat-inactivated foetal bovine serum (FBS; Gibco: 10500064) in a humidified incubator at 37°C with 5% CO2. THP-1 monocytes were differentiated into macrophages in RPMI + 10 mM HEPES + 10% FBS + 10 ng/mL phorbol 12-myristate 13-acetate (PMA; Sigma) and seeded at 1×10^5^ cells per well of a 96-well tissue culture dish and further treated with 66.8 μM concentration of bifidococcin664 separately, before incubating for 15 hours. Overnight cultures of bacteria were diluted 1:100 into fresh LB broth and grown until mid-exponential phase. Bacteria were then washed twice with PBS and diluted to 1×10^7^ cfu/mL in RPMI + 10 mM HEPES + 10% FBS and 100 μl of bacteria was added to each well that contain bacteriocin treated macrophages. Plates were then centrifuged at 300 *g* for 5 min to synchronize infections. Bacteria–macrophage co-culture was incubated at 37°C/5% CO2 for 30 mins to allow for phagocytosis. Cells were then washed three times in PBS and medium was replaced with above culture medium supplemented with 300 μg/mL gentamicin to eliminate extracellular bacteria. Cell were again incubated at 37°C/5% CO2 for 1 hour. Cells were then washed three times with PBS and medium for cells for later time points was replaced with culture medium supplemented with 300 μg/mL gentamicin and incubated for a further 3 hours. Intracellular bacterial load was enumerated by lysing macrophages in PBS + 1% Triton X-100 for 10 mins at room temperature, serially diluting cultures and plating on LB agar. Plates were incubated overnight at 37°C and colonies counted the following day.

### Cytokine analysis and Caco2 cells treatment

The Caco2 cells purchased from Public Health England (ECACC catalogue no. 09042001). were cultured in high glucose Dulbecco’s modified Eagle medium supplemented with 10% foetal bovine serum (FBS), 2 mM L-Gln, non-essential amino acids, 100 U/mL penicillin/100 μg/mL streptomycin (Gibco, Waltham, MA, USA), and 20 mM HEPES. The cells (P14) were seeded at 2×10^4^ per well in 96-well and waited to attain confluence prior to Bifidococcin_664 treatments (17.368 μM, 43.402 μM, and 86.8403) in 200μL of culture media per well and incubated for 24hours. The supernatant was collected and stored for cytokine analysis.

Supernatant collected after each treatment of the cells was stored at −80°C. IL-6, IL-8, IL-15 and TSLP concentrations of the supernatants were measured by using U-plex Meso Scale Discovery platforms (MSD Inc., Rockville USA). The assay was performed as per the manufacturer’s instructions and plates were read using a MESO QuickPlex SQ 120 instrument. The cytokine concentrations were calculated by MSD Discovery workbench software based on the standard curves of calibrators provided by the MSD kit. The estimation of TNF-α, the supernatant was performed using commercially available kits for cytokine detection (ELISA DUO, R&D systems, USA). Preparation of reagents, standards, and protocol were followed according to the manufacturer’s instructions. The absorbance was read using ELISA reader (BIO-RAD). The detection ranges for TNF-α, 15.6 pg/ml− 1000 pg/ml.

### Statistics

Statistical analyses were performed using analysis of variance (ANOVA) in the GraphPad Prism software (version 8 USA). All experiments performed in duplicates and triplicate and error bar indicates standard deviation. Significance was defined as *P*<0.05.

---

**Supplementary Figure 1:**
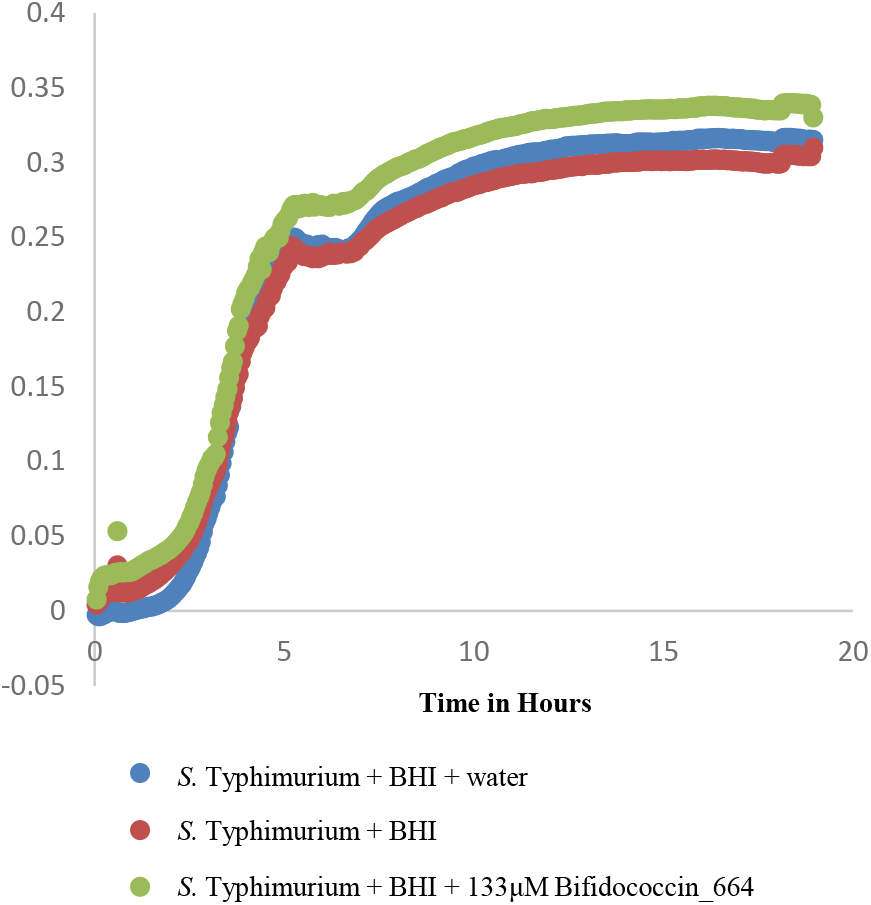
Growth curve of *S*. Typhimurium 3580 in BHI showing resistance to tested bacteriocin concentration. Growth followed using OD_600_.

